# Use of the Azure Kinect to measure foot clearance during obstacle crossing

**DOI:** 10.1101/2021.08.16.455774

**Authors:** Kohei Yoshimoto, Masahiro Shinya

**Author notes:** Corresponding author: Masahiro Shinya, Hiroshima University, 1-7-1 Kagamiyama, Higashi-Hiroshima, 739-8521, Japan.

## Abstract

Obstacle crossing is a typical adaptive locomotion known to be related to the risk of falls. Previous conventional studies have used elaborate and costly optical motion capture systems, which not only represent a considerable expense but also require participants to visit a laboratory. To overcome these shortcomings, we aimed to develop a practical and inexpensive solution for measuring obstacle-crossing behavior by using the Microsoft Azure Kinect, one of the most promising markerless motion capture systems. We validated the Azure Kinect as a tool to measure foot clearance and compared its performance to that of an optical motion capture system (Qualisys). We also determined the effect of the Kinect sensor placement on measurement performance. Sixteen healthy young men crossed obstacles of different heights (50, 150, and 250 mm). Kinect sensors were placed in front of and beside the obstacle as well as diagonally between those positions. As indices of measurement quality, we counted the number of measurement failures and calculated the systematic and random errors between the foot clearance measured by the Kinect and Qualisys. We also calculated the Pearson correlation coefficients between the Kinect and Qualisys measurements. The number of measurement failures and the systematic and random error were minimized when the Kinect was placed diagonally in front of the obstacle on the same side as the trail limb. The high correlation coefficient (r > 0.890) observed between the Kinect and Qualisys measurements suggests that the Azure Kinect has excellent potential for measuring foot clearance during obstacle-crossing tasks.

## Introduction

Falls may have a serious impact on health, independence, and quality of life in the elderly population. Among elderly people over 75 years old, 24% of those who fell were severely injured, and 6% suffered fractures (Tinetti et al., 1988). Because approximately 30% of falls among elderly people are due to contact with obstacles (Berg et al., 1997; Overstall et al., 1977), understanding the mechanisms behind obstacle-crossing behavior in the interest of fall prevention is essential in an aging modern society.

Quantitative evaluation of gait parameters during obstacle negotiation, such as clearance (the vertical distance from the foot to the obstacle), is crucial in obstacle-crossing studies. Evidence from previous studies implies that the clearance would be an index that could potentially predict the risk of contact with obstacles. There is evidence that the age-related decline in physical function affects the kinematics of obstacle crossing, for example, by increasing clearance asymmetry (Fabio et al., 2004; Pan et al., 2016; Tomar and Gupta, 2012). Elderly people are sometimes not aware of the decline in their physical function, potentially increasing their risk of falls (Sakurai et al., 2013). Another previous study in healthy adults reported that greater variability in clearance is associated with a greater risk of contact with obstacles (Rietdyk and Rhea, 2011).

Despite the potential practicality, obstacle-crossing behavior and foot clearance may vary between populations and may be influenced by various conditions because obstacle crossing is a complex and highly coordinated locomotor task. The effect of aging on the absolute value of foot clearance is controversial (reviewed in (Galna et al., 2009): some studies reported no effect of aging on foot clearance (Chen et al., 1991; Lowrey et al., 2007), whereas others reported that healthy elderly adults showed larger (Lu et al., 2006; Muir et al., 2019; Yen et al., 2009) or smaller clearance (Maidan et al., 2018; McFadyen and Prince, 2002) than young adults. Foot clearance is also known to be affected by dual-tasking conditions (Harley et al., 2009), trial repetition (Heijnen et al., 2012), and interlimb interaction (Miura and Shinya, 2021). Therefore, to better understand the mechanisms of motor control in obstacle crossing, further research is needed in various populations and various contexts.

The most significant barriers to obstacle-crossing research include the high cost and lack of portability of optical motion capture systems. Markerless motion capture is a promising solution to overcome the weakness of optical motion capture systems that require reflecting markers. The Microsoft Kinect is a low-cost, portable device containing a body tracking system that can measure three-dimensional joint positions. The usefulness of the Kinect body tracking system has been validated in gait and posture studies and clinical settings (reviewed in Clark et al., 2019). In contrast to most gait and posture tasks such as regular walking and standing, measuring foot clearance during obstacle crossing tasks requires data on the position of an external object (i.e., the obstacle) in addition to the subject’s skeleton. This means that the body skeleton data measured by the Kinect, which are in camera-centered coordinates, should be transformed into the global coordinate system that defines the location of the obstacle. Although one study used the Kinect v2 to measure foot clearance (Maidan et al., 2018), coordinate transformation of Kinect data has not yet been validated.

The effect of the location of the Kinect sensor also needs to be investigated. In gait analysis, researchers often placed the first- and second-generation Kinect sensors in front of the participant (Eltoukhy et al., 2017; Kharazi et al., 2015; Mentiplay et al., 2015; Tanaka et al., 2018; Xu et al., 2015), while a few studies placed the sensor diagonally in front of the participant (Pfister et al., 2014; Yeung et al., 2021). A recent study showed that the optimal camera angle might differ depending on the version of Kinect sensor that is used (Yeung et al., 2021). Yeung et al. (2021) recorded the lower limb joint angles during treadmill walking with the Kinect v2 and the Azure Kinect at five camera angles: 0°, 22.5°, 45°, 67.5°, and 90°. The results showed that the error was minimized at oblique rather than head-on camera angles. In this study, we aimed to find the best location for the Azure Kinect sensor to measure obstacle-crossing behavior.

Since self-occlusion drastically impacts measurement quality, the difference in the measurement of the foot clearance between the lead and trail limbs should be tested. Additionally, given that the Kinect estimates the location of body landmarks with a machine learning-based algorithm (Wang et al., 2015), the effect of different kinematics on obstacle crossing should also be tested. It is known that lower limb kinematics differ between the lead and trail limbs (Chou and Draganich, 1997; Lu et al., 2006) and between different obstacle heights (Austin et al., 1999).

The purpose of this study was to 1) validate the Azure Kinect for assessing foot clearance during obstacle crossing and 2) determine the effects of the Kinect sensor location on the measurement. To this end, obstacle-crossing behavior for obstacles of different heights was recorded by Azure Kinect sensors from four different viewing angles. Then, we compared the foot clearances of the lead and trail limbs between the Kinect and conventional optical motion capture systems. To identify the best Kinect location, we compared the quality of measurements between the different Kinect locations.

## Methods

Sixteen healthy young men (age: 21.6 ± 0.8 years; height: 170.7 ± 6.2 cm; weight: 59.4 ± 6.6 kg) participated in this study. The participants did not have any disability that could influence walking. The purpose and precautions of the study were fully explained to the participants in advance, and informed consent was obtained. The study was conducted in accordance with the Declaration of Helsinki and approved by the local ethics committee (approval number: 01-31).

Participants walked barefoot at their own pace and crossed an obstacle. They were instructed to step over the obstacle with their right foot at the 7th step and then continue walking at least four more steps (Fig. 1a). Three obstacle heights were tested: 50, 150, and 250 mm. The obstacle was a 1.00 m long plastic tube with a diameter of 10 mm.

**Fig. 1.**
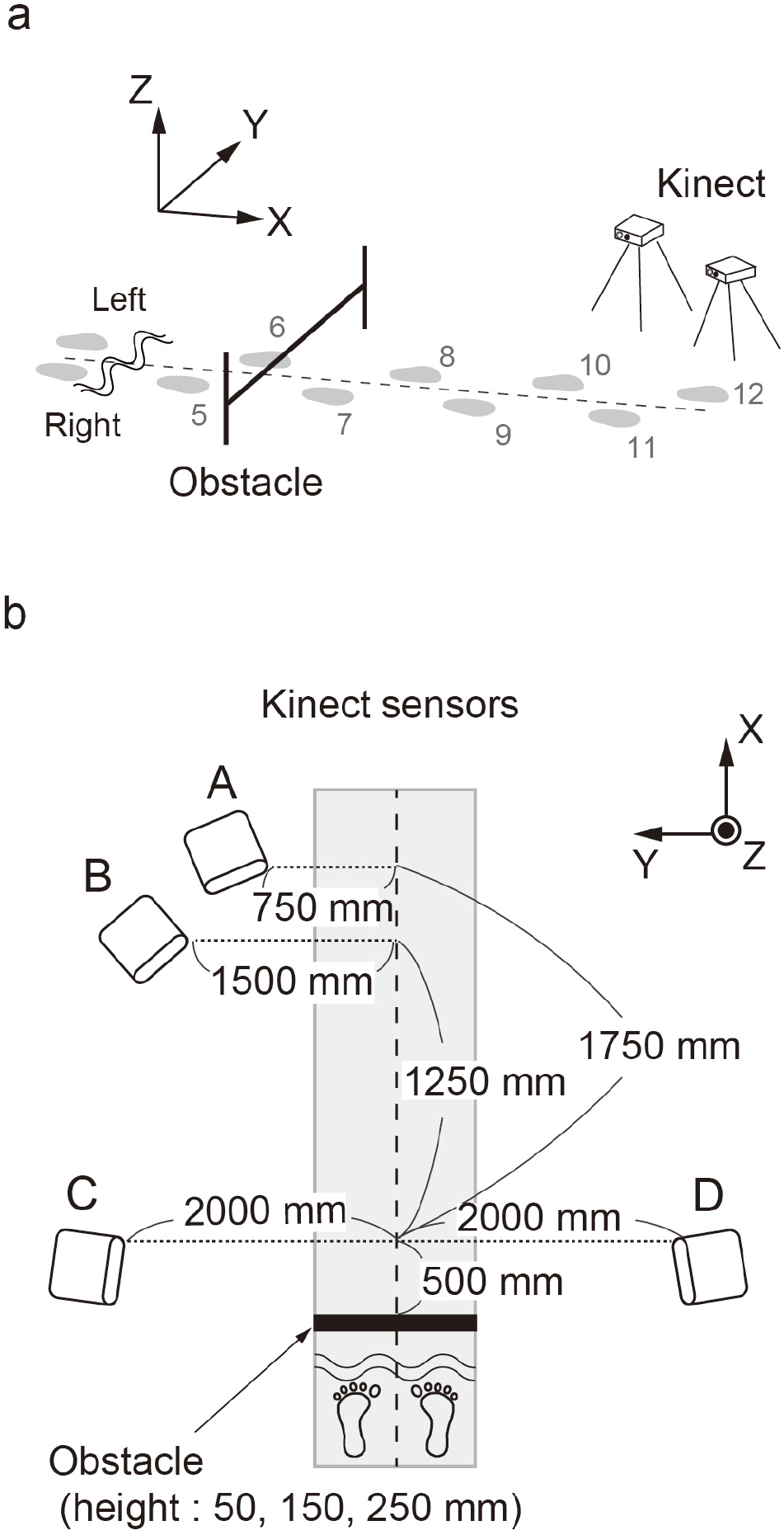
Experimental diagram. Participants crossed an obstacle (height: 50, 150, 250 mm) at the 7th step, with their right limb being the lead limb (a). The Kinect sensors were set up in two different layouts (b). In one layout, the sensor positions (location A and location B) were located at a 20° angle and 40° angle relative to the walkway on the trail limb side; in the other layout, the sensor positions (location C and location D) were located on opposite sides, at a 75° angle and a -75° angle.

Whole-body kinematics was measured by two Azure Kinect sensors (Microsoft) and eight optical cameras (Miqus M3, Qualisys, Sweden). In this study, the Kinect sensors were set up in two different layouts. In one layout, two Kinect sensors were placed at 20° and 40° diagonal positions on the trail limb side (location A and location B in Fig 1b). In another layout, Kinect sensors were placed at 75° and -75° angles, and images of the walking participants were captured from both the lead and trail limb sides (location C and location D in Fig 1b). Ten trials were recorded for each obstacle height and Kinect layout. Any trial in which the participant hit the obstacle was considered a failed trial, and the participant was allowed to repeat the trial until ten successful trials were recorded. The Azure Kinect Body Tracking SDK provided 3D skeleton data constructed of 32 body points at a sampling rate of 30 Hz.

In our preliminary recordings and a previous study (Naeemabadi et al., 2018), it was observed that infrared markers placed on the ankle and knee for optical motion capture could hinder the recording of depth images by the Azure Kinect sensor. Instead of placing a marker on the knee and ankle, we created a rigid body model for the shank segment before the recording session. The rigid body model was created from markers attached to the medial and lateral femoral epicondyles and on the medial and lateral malleoli as well as two markers on the middle of the shank segment. When we recorded the obstacle-crossing task using the Kinect and Qualisys, we removed the markers on the medial or lateral femoral epicondyle and the medial or lateral malleolus, and the locations of these points were estimated from the other four markers of the shank rigid body (Fig. 2). Except for the shank segments, significant interference of the infrared reflective markers with Qualisys recording was not observed. We placed markers on the left and right tragus, acromion, anterior superior iliac spine (ASIS), posterior superior iliac spine (PSIS), greater trochanter (GTR), first metatarsal bone, and radial and ulnar styloid processes.

**Fig. 2.**
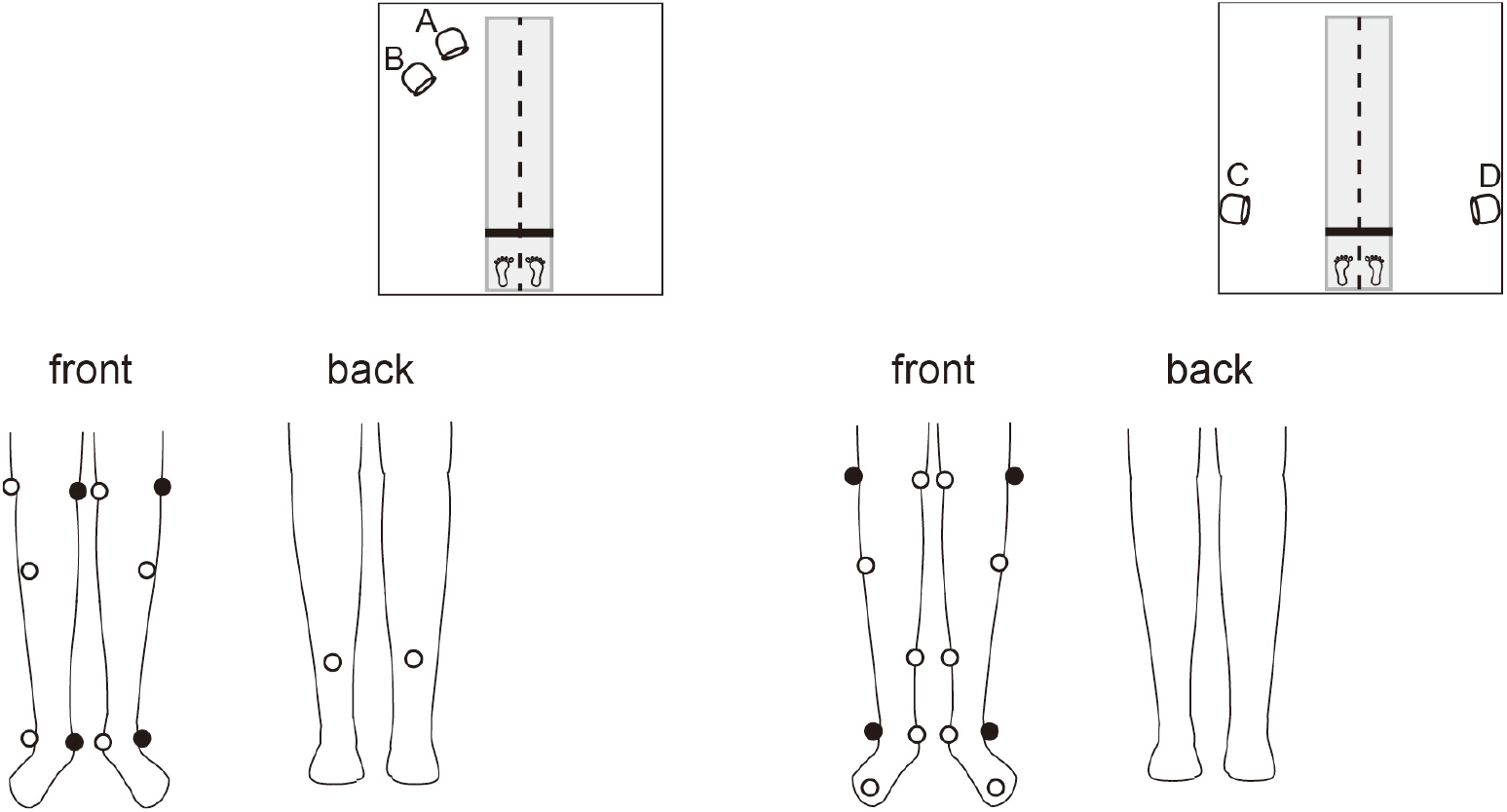
Reflective marker placement. A rigid body model was made for the shank segment from six infrared reflective markers. During the recording obstacle-crossing behavior, the markers on the side of the knee and ankle facing the Kinect were removed to minimize the interference between the Azure Kinect and the optical motion capture.

The original skeleton data obtained from the Kinect SDK were in a coordinate system based on the depth camera of the Kinect. The skeleton data were transformed into the laboratory coordinate system for calculation of the foot clearance and the distance between the foot and obstacle. For this transformation, we carried out the following steps (Fig. 3). 1) We placed two boards in the laboratory, and the point cloud data (resolution: 640×576) were collected from the Kinect depth image. 2) The plane of the floor was estimated using the Random Sample Consensus (RANSAC) algorithm in the MATLAB computer vision toolbox, and the tilt angle of the Kinect sensor was calculated. 3) After correction for the tilt angle of the Kinect, the distance from the Kinect camera to the floor plane was calculated as the z coordinate of the center of gravity of the floor plane. 4) The plane of one of the two boards was estimated, and its tilt angle was calculated. 5) After correction for the tilt angle of the board plane, the coordinates of the intersection point on the line extending through the two boards were calculated. Once the transformation matrix from Kinect coordinates to laboratory coordinates was obtained, the skeleton data were transformed into laboratory coordinates.

**Fig. 3.**
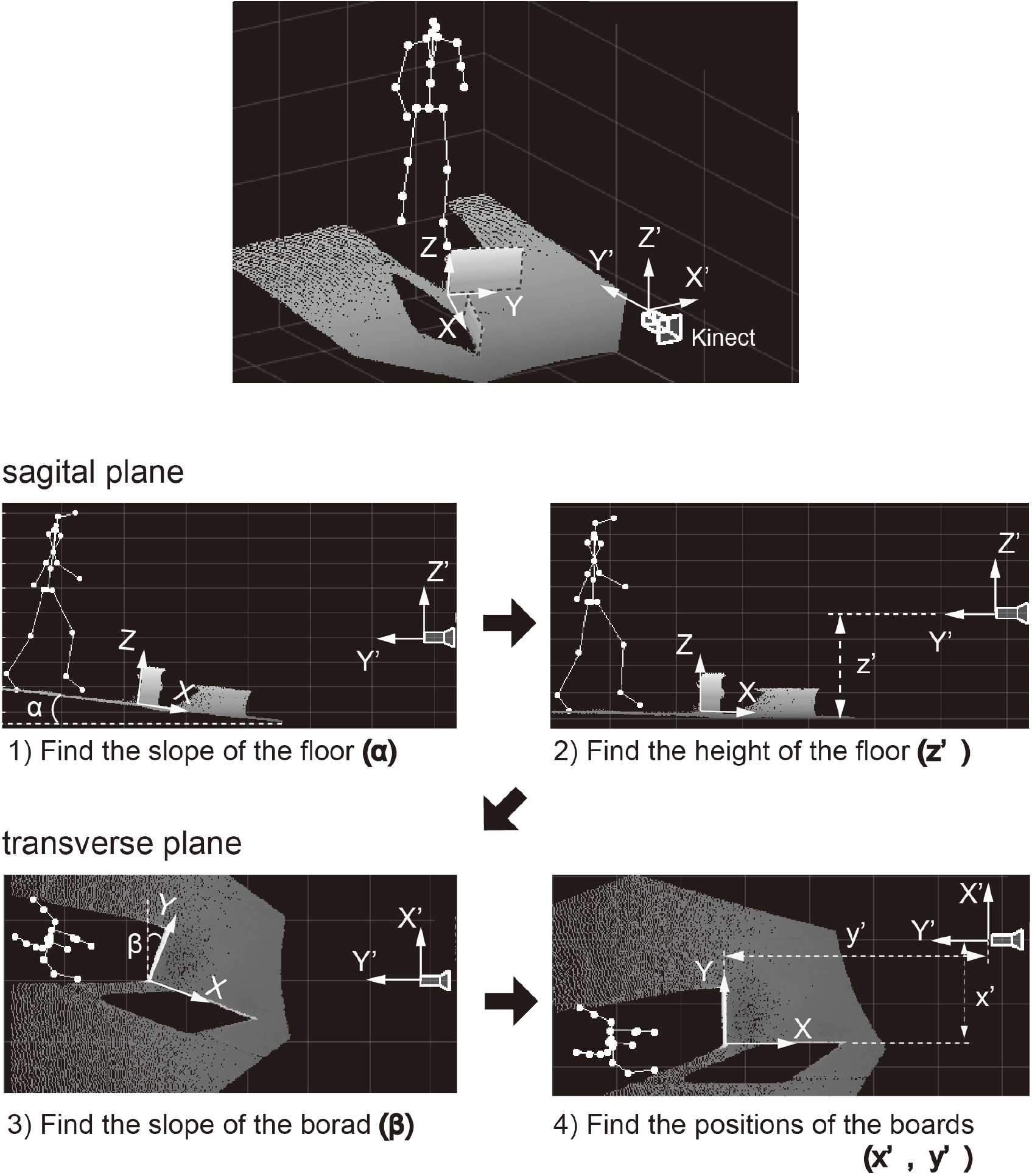
Transformation from the Kinect’s skeleton coordinates (X′, Y′, Z′) to laboratory coordinates (X, Y, Z). Point cloud (resolution: 640×576) data were used for the transformation.

Kinect measurements with a confidence level of high or medium were used, and those with low or no confidence were replaced with estimated values calculated by spline interpolation. Although the data with low or no confidence were discarded, apparently incorrect data were observed where the position of the toe marker was estimated to be external to the body. These trials were defined by multiple peaks with a significant estimated dropdown greater than 100 mm in the vertical displacement of the toe during obstacle crossing. Trials in which the foot trajectory was estimated to be lower than the obstacle were also regarded as erroneous trials. These errors in the Kinect measurements were considered measurement failures. We counted the number of measurement failures for each Kinect location and used these counts as an index of measurement quality.

The Kinect data and the Qualisys data were smoothed using a zero-lag second-order low-pass digital Butterworth filter with a cutoff frequency of 5 Hz. Spline interpolation was used to resample the Kinect data to 240 Hz from 30 Hz. Foot clearance was defined as the vertical distance from the toe to the obstacle when the toe was directly above the obstacle. The clearance calculated by the Qualisys was compensated by subtracting the vertical offset of the toe marker during standing. We also calculated residuals in the clearance by subtracting the Qualisys data from the Kinect data. Then, we defined the systematic error as the within-subject mean value of the residuals and the random error as the within-subject standard deviation of the residuals (Fig. 4).

**Fig. 4.**
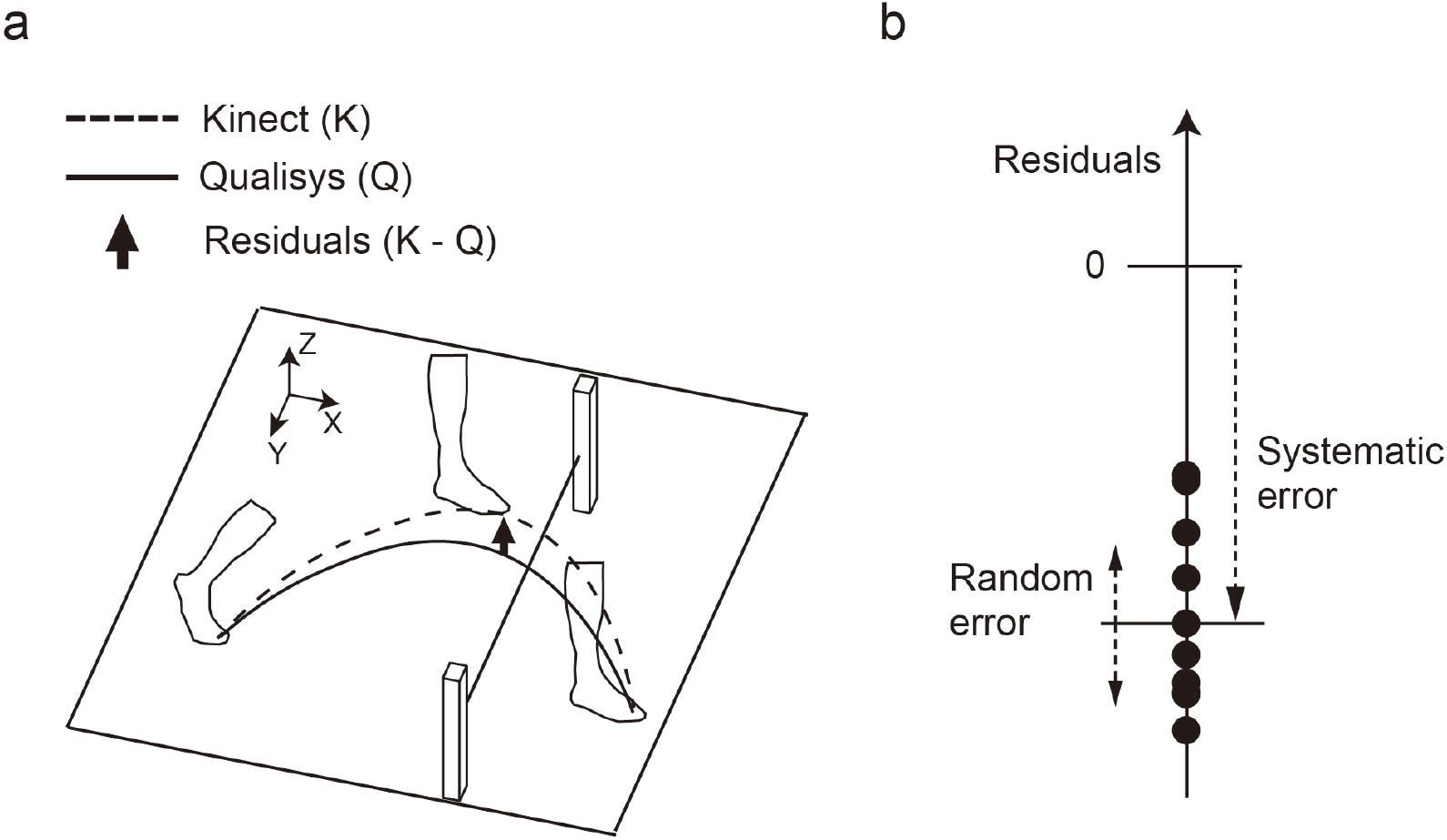
Calculation of the systematic and random error. Residuals were calculated by subtracting Qualisys from Kinect (a). The systematic error and random error were defined as the mean and standard deviation, respectively, of the residuals (b).

Friedman tests were used to evaluate differences in the number of measurement failures, the systematic error, and the random error among Kinect locations. Friedman tests were performed separately for the lead limb in the 50 mm obstacle condition (L50), for the lead limb in the 150 mm obstacle height condition (L150), for the lead limb in the 250 mm obstacle height condition (L250), for the trail limb in the 50 mm obstacle condition (T50), for the trail limb in the 150 mm obstacle height condition (T150), and for the trail limb in the 250 mm obstacle height condition (T250). The significance level of 0.05 was used for the Friedman tests. Kendall’s W was used as an index of effect size for the Friedman tests. If a significant main effect was observed in the Friedman tests, multiple comparisons were performed with Conover’s test, and the significance level was adjusted by using the Holm correction (6 pairs). For each limb and obstacle height, within-subject averages were calculated in the foot clearance, and systematic and random errors were calculated in the foot clearance if the number of measurement failures was 5 or less. If the number of measurement failures exceeded 5 (out of 10 trials), the data were excluded from the subsequent analyses. For these variables, median [minimum, maximum] was used as descriptive. A one-sample Wilcoxon signed-rank test was performed to determine whether the systematic error was significantly different from 0. Pearson’s correlation analysis was performed between the within-subject average of the clearances measured by Kinect and that measured by Qualisys. As location B (i.e., oblique viewing angle) showed the best measurement performance (see Results for details), the systematic error and the random error were compared between limbs (lead, trail) and obstacle heights (50, 150, 250 mm) by using pairwise Wilcoxon signed-rank tests with a significance level adjusted by using the Holm correction (15 pairs).

## Results

Friedman tests revealed that the number of measurement failures was different depending on the Kinect location except for the L50 measurement (Table 1). According to Conover’s multiple comparison tests, significantly fewer measurement failures occurred in location B than in locations C and D for the L150 measurement. Locations A and B showed significantly fewer measurement failures than location C for the L250 measurement. In T50 and T150, locations A, B, and C showed significantly fewer measurement failures than location D. In T250, location C showed significantly fewer measurement failures than locations A and D.

**Table 1.**
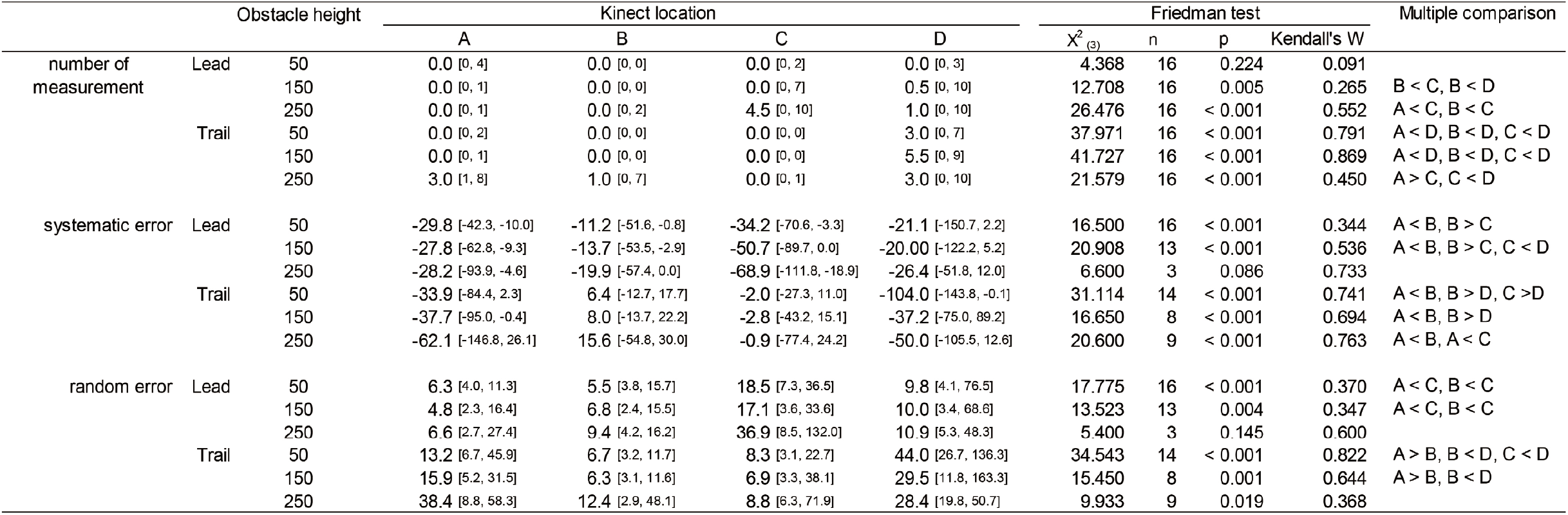
Comparison of foot clearance measurement performance between Kinect locations. Median, maximum, and minimum values were used as descriptive statistics for the number of measurement failures, systematic error, and random error.

The systematic error of the foot clearance was also influenced by the Kinect location. Friedman tests showed significant main effects of the Kinect location for L50, L150, T50, T150, and T250. There was no significant main effect observed for L250 because the degree of freedom for the test was very small due to the large number of measurement failures. Given that the zero systematic error is the best performance, the Kinect measurement from location B showed a greater agreement with the Qualisys measurement than the measurements from locations A and C for L50. In the L150 condition, there were significant differences in the systematic error between the Kinect location A and B, between B and C, and between C and D. In the T50, significant differences were observed between location A and B, between B and D, and between C and D. In the T150, there were significant differences between location A and B, and between B and D. In the T250, there were significant differences between location A and B, and between A and C. A one-sample Wilcoxon signed-rank test revealed that the systematic error of the foot clearance in the lead limb was significantly less than 0 for all the Kinect locations and obstacle heights tested (Table 2). The systematic error of the foot clearance in the trail limb in the measurement from locations B and C was not significantly different from 0 except for the 150 mm obstacle condition measured from location B. The systematic error of the foot clearance in the trail limb in the measurement from locations A and D was significantly different from 0 except for the 150 mm obstacle condition measured from location D.

**Table 2.** Descriptive statistics (median, maximum, and minimum) and results of one-sample Wilcoxon signed-rank tests (null hypothesis: = 0) for the systematic error.

|  | Obstacle height | location A |  | location B |  | location C |  | location D |  |
| --- | --- | --- | --- | --- | --- | --- | --- | --- | --- |
| systematic error (Lead) | 50 | -29.8 [-42.3, -10.0] | p < 0.001 | -11.2 [-51.6, -0.8] | p < 0.001 | -34.2 [-70.6, -3.3] | p < 0.001 | -21.1 [-150.7, 2.2] | p < 0.001 |
|  | 150 | -27.8 [-62.8, -9.3] | p < 0.001 | -13.7 [-53.5, -2.9] | p < 0.001 | -50.7 [-89.7, 0.0] | p < 0.001 | -20.0 [-122.2, 5.2] | p < 0.001 |
|  | 250 | -28.2 [-93.9, -4.6] | p < 0.001 | -19.9 [-57.4, 0.0] | p < 0.001 | -68.9 [-111.8, -18.9] | p = 0.004 | -26.4 [-51.8, 12.0] | p = 0.027 |
| systematic error (Trail) | 50 | -33.9 [-84.4, 2.3] | p < 0.001 | 6.4 [-12.7, 17.7] | p = 0.105 | -2.0 [-27.3, 11.0] | p = 0.404 | -104.0 [-143.8, -0.1] | p < 0.001 |
|  | 150 | -37.7 [-95.0, -0.4] | p < 0.001 | 8.0 [-13.7, 22.2] | p = 0.004 | -2.8 [-43.2, 15.1] | p = 0.404 | -37.2 [-75.0, 89.2] | p = 0.195 |
|  | 250 | -62.1 [-146.8, 26.1] | p < 0.001 | 15.6 [-54.8, 30.0] | p = 0.303 | -0.9 [-77.4, 24.2] | p = 0.782 | -50.0 [-105.5, 12.6] | p = 0.020 |

The alternative hypothesis specifies that the median is different from 0.

For the random error, Friedman tests showed significant main effects of the Kinect location for L50, L150, T50, T150, and T250. There was no significant main effect observed for L250. In the L50 and L150 conditions, significantly smaller random errors were observed in locations A and B than in location C. In T50, significant differences were observed between locations A and B, between B and D, and between C and D. In the T150 condition, significantly smaller random error was observed in location B than in location A and D.

Correlations between the foot clearances measured by the Kinect and Qualisys are illustrated in Fig. 5. Overall, the Kinect measurements from location B showed good agreement with the Qualisys measurements for both the lead and trail limbs (r > 0.890).

**Fig. 5.**
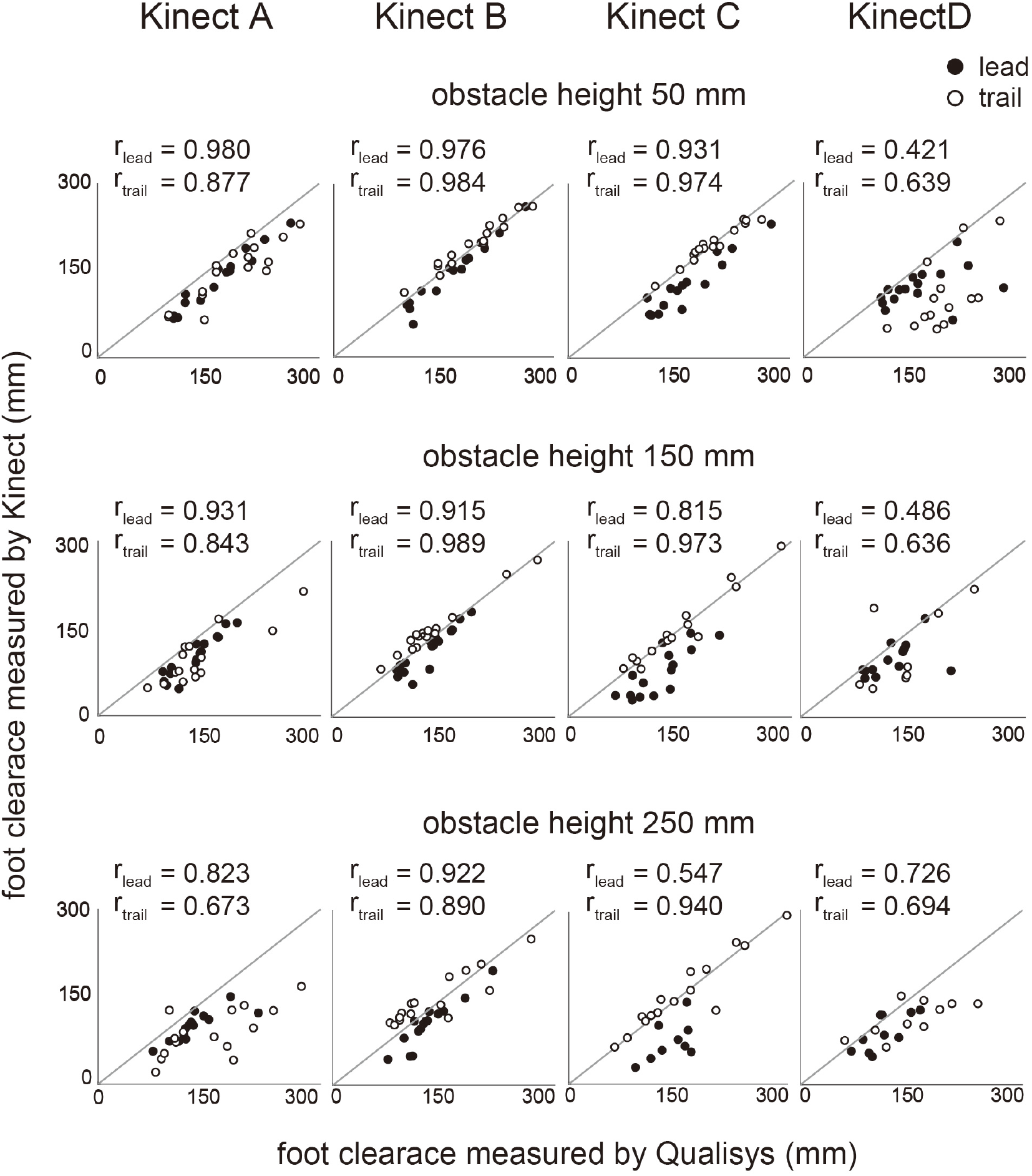
Relationship between the foot clearance measured by the Kinect and Qualisys. The diagonal straight lines represent the identity line, y = x. The black circles represent the data for the lead limb, and the white circles represent those for the trail limb. Pearson’s correlation coefficients were calculated separately for the lead and trail limbs.

The systematic and random error of the foot clearance in the measurement from location B were compared between limbs and obstacle height conditions (Table 3). Significantly smaller systematic errors were observed in the lead limb than in the trail limb. In the comparisons between obstacle height conditions, there was a significant difference in the systematic error between L50 and L150, and between L150 and L250. The random error in L50 was significantly smaller than that in T250.

**Table 3.** Systematic error and random error measured in location B were compared between limbs and obstacle height conditions by using Wilcoxon signed-rank tests. Median, maximum, and minimum values are shown as descriptive.

|  | Obstacle height | Lead | Trail |  |
| --- | --- | --- | --- | --- |
| systematic error | 50 | -11.2 [-51.6, -0.8] * | 6.4 [-12.7, 17.7] | # |
|  | 150 | -13.7 [-53.5, -2.9] * | 8.0 [-13.7, 22.2] | # |
|  | 250 | -19.9 [-57.4, 0.0] | 15.6 [-54.8, 30.0] | # |
| random error | 50 | 5.5 [3.8, 15.7] | 6.7 [3.2, 11.7] | * |
|  | 150 | 6.8 [2.4, 15.5] | 6.3 [3.1, 11.6] | * |
|  | 250 | 9.4 [4.2, 16.2] | 12.4 [2.9, 48.1] |  |

Holm correction's significant difference \*: $p < 0.05$ (v.s. 250 mm obstacle height); #: $p < 0.05$ (Lead v.s. Trail)

## Discussion

In this study, we validated Azure Kinect sensors for the measurement of foot clearance during an obstacle-crossing task. The results suggest that foot clearance was reliably measured by the Azure Kinect sensor and that the sensor should be placed diagonally in front of the obstacle on the same side as the trail limb (location B in this study), which is consistent with a previous study (Yeung et al., 2021). The number of measurement failures was larger when the Kinect was placed by the side of the obstacle (locations C and D) than when it was placed directly or diagonally in front (locations A and B). Contralateral measurements (i.e., the lead limb measured from location C and the trail limb measured from location D) had more measurement failures, larger systematic error, and larger random error than ipsilateral measurements, perhaps because the closer occluded the farther limb (Pfister et al., 2014; Seo et al., 2016) (typical depth images are shown in supplementary Fig. S1). Notably, the sensors that were placed beside the obstacle did not perform well even for ipsilateral measurements (typical depth image shown in supplementary Fig. S2).

Based on the systematic and random error, Kinect location B showed better performance than any other tested placement. Correlation analysis also suggested that the foot clearance measured by a single diagonally placed Azure Kinect was reliable for both the lead and trail limbs. The observed poor performance of the sagittally placed Kinects might be due to self-occlusion, as discussed in the previous paragraph. The position in front of the walking subject (location A) was also not the optimal sensor location. The systematic and random error were larger for location A than for location B. This might be because the oblique camera view angle provides information about both the sagittal and frontal planes (Yeung et al., 2021).

The poor measurement performance of the Kinect in front of the subject was significant for the trail limb crossing the high (250 mm) obstacle. This was probably because the trail limb, unlike the lead limb, crossed over the high obstacle by flexing the knee joint rather than the hip and ankle joints (Chou and Draganich, 1997). This knee flexion hides the foot segment behind the thigh segment, leading to unreliable estimation of the toe position during obstacle crossing.

As discussed above, the Azure Kinect can be a reliable portable solution for measuring foot clearance during obstacle-crossing tasks. Previous studies with older versions of the Kinect reported that toe markers generated a large amount of noise (Wang et al., 2015; Xu et al., 2015). In contrast, a recent study reported that the Azure Kinect had improved toe tracking performance compared to the Kinect v2 (Albert et al., 2020). Additionally, the depth sensor of the Azure Connect has a higher resolution (640×576) than that of the Kinect v2 (512×424), which may contribute to the reduction of the error in toe positioning.

Caution should be taken in drawing within-subject comparisons of foot clearance between the lead and trail limbs or between different obstacle height conditions. Our results showed that the diagonally placed Azure Kinect did not overestimate or underestimate the foot clearance of the trail limb; in contrast, the foot clearance of the lead limb was underestimated by approximately 10 to 20 mm. In the 250 mm obstacle height condition, the systematic error of the foot clearance for the lead limb was different from that in the 50 and 150 mm obstacle height conditions, and the random error for the trail limb was greatest at this obstacle height. These differences in systematic and random error between lead and trail limbs or obstacle heights may be due to the differences in posture between conditions. This might be because the machine learning-based posture estimation algorithm of the Azure Kinect system uses information from the whole body to estimate where the centers of the joints (ankle, foot, etc.) are positioned. Similarly, the kinematics of obstacle crossing kinematics may differ between healthy young people and elderly people or patients with various diseases (Chen et al., 2019). Therefore, future studies should validate whether the Azure Kinect can measure foot clearance in those populations.

## Acknowledgments

This study was supported by JSPS KAKENHI Grant Number 17H04750.

## Supplementary Material

**Fig. S1.**
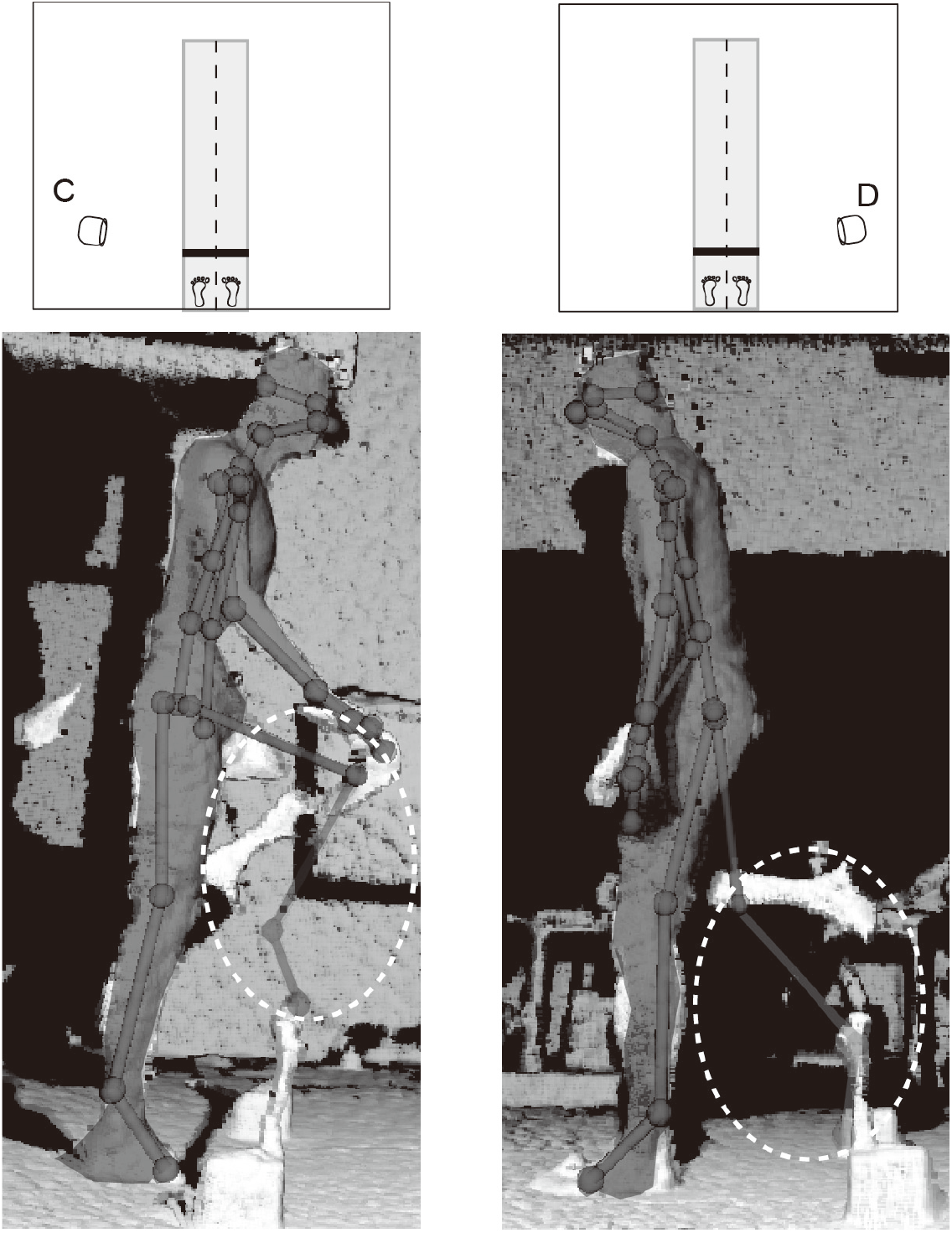

**Fig. S2.**
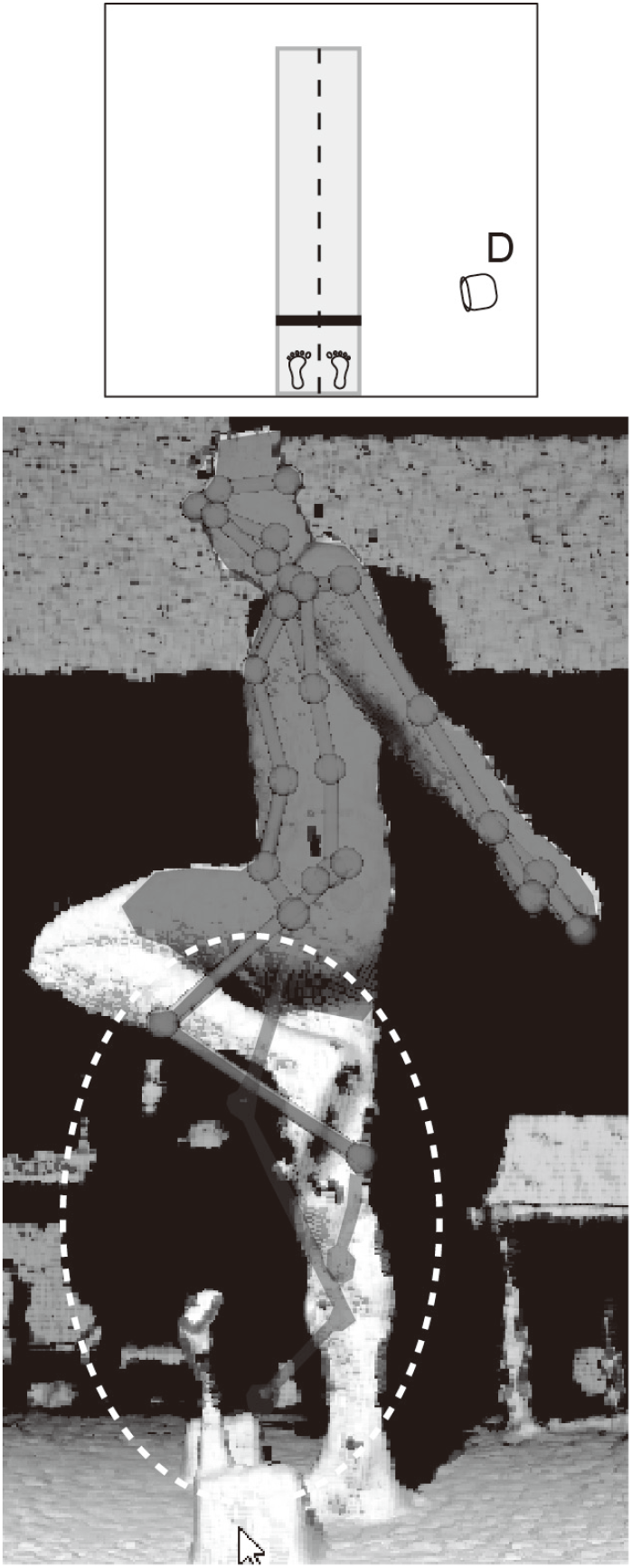

## References

Albert, J.A., Owolabi, V., Gebel, A., Brahms, C.M., Granacher, U., Arnrich, B., 2020. Evaluation of the pose tracking performance of the azure kinect and kinect v2 for gait analysis in comparison with a gold standard: A pilot study. Sensors (Switzerland) 20, 1–22.

Austin, G.P., Garrett, G.E., Bohannon, R.W., 1999. Kinematic analysis of obstacle clearance during locomotion. Gait Posture 10, 109–120.

Berg, W.P., Alessio, H.M., Mills, E.M., Tong, C., 1997. Circumstances and consequences of falls in independent community-dwelling older adults. Age Ageing 26, 261–268.

Chen, H.C., Ashton-Miller, J.A., Alexander, N.B., Schultz, A.B., 1991. Stepping over obstacles: Gait patterns of healthy young and old adults. Journals Gerontol. 46, 196–203.

Chen, N., Xiao, X., Hu, H., Chen, Y., Song, R., Li, L., 2019. Identify the alteration of balance control and risk of falling in stroke survivors during obstacle crossing based on kinematic analysis. Front. Neurol. 10, 1–11.

Chou, L.S., Draganich, L.F., 1997. Stepping over an obstacle increases the motions and moments of the joints of the trailing limb in young adults. J. Biomech. 30, 331–337.

Clark, R.A., Mentiplay, B.F., Hough, E., Pua, Y.H., 2019. Three-dimensional cameras and skeleton pose tracking for physical function assessment: A review of uses, validity, current developments and Kinect alternatives. Gait Posture 68, 193–200.

Eltoukhy, M., Oh, J., Kuenze, C., Signorile, J., 2017. Improved kinect-based spatiotemporal and kinematic treadmill gait assessment. Gait Posture 51, 77–83.

Fabio, R.P. Di, Kurszewski, W.M., Jorgenson, E.E., Kunz, R.C., 2004. Footlift Asymmetry During Obstacle Avoidance in High-Risk Elderly. J. Am. Geriatr. Soc. 2088–2093.

Galna, B., Peters, A., Murphy, A.T., Morris, M.E., 2009. Obstacle crossing deficits in older adults:A systematic review. Gait Posture 30, 270–275.

Harley, C., Wilkie, R.M., Wann, J.P., 2009. Stepping over obstacles: Attention demands and aging. Gait Posture 29, 428–432.

Heijnen, M.J.H., Muir, B.C., Rietdyk, S., 2012. Factors leading to obstacle contact during adaptive locomotion. Exp. Brain Res. 223, 219–231.

Kharazi, M.R., Memari, A.H., Shahrokhi, A., Nabavi, H., Khorami, S., Rasooli, A.H., Barnamei, H.R., Jamshidian, A.R., Mirbagheri, M.M., 2016. Validity of microsoft kinectTM for measuring gait parameters. 2015 22nd Iran. Conf. Biomed. Eng. ICBME 2015 375–379.

Lowrey, C.R., Watson, A., Vallis, L.A., 2007. Age-related changes in avoidance strategies when negotiating single and multiple obstacles. Exp. Brain Res. 182, 289–299.

Lu, T.W., Chen, H.L., Chen, S.C., 2006. Comparisons of the lower limb kinematics between young and older adults when crossing obstacles of different heights. Gait Posture 23, 471–479.

Maidan, I., Eyal, S., Kurz, I., Geffen, N., Gazit, E., Ravid, L., Giladi, N., Mirelman, A., Hausdorff, J.M., 2018. Age-associated changes in obstacle negotiation strategies: Does size and timing matter? Gait Posture 59, 242–247.

McFadyen, B.J., Prince, F., 2002. Avoidance and accommodation of surface height changes by healthy, community-dwelling, young, and elderly men. Journals Gerontol. - Ser. A Biol. Sci. Med. Sci. 57, B166–B174.

Mentiplay, B.F., Perraton, L.G., Bower, K.J., Pua, Y.H., McGaw, R., Heywood, S., Clark, R.A., 2015. Gait assessment using the Microsoft Xbox One Kinect: Concurrent validity and inter-day reliability of spatiotemporal and kinematic variables. J. Biomech. 48, 2166–2170.

Miura, Y., Shinya, M., 2021. Foot clearance when crossing obstacles of different heights with the lead and trail limbs. Gait Posture 88, 155–160.

Muir, B.C., Haddad, J.M., van Emmerik, R.E.A., Rietdyk, S., 2019. Changes in the control of obstacle crossing in middle age become evident as gait task difficulty increases. Gait Posture 70, 254–259.

Naeemabadi, Mr., Dinesen, B., Andersen, O.K., Hansen, J., 2018. Investigating the impact of a motion capture system on Microsoft Kinect v2 recordings: A caution for using the technologies together. PLoS One 13.

Overstall, P.W., Exton-Smith, A.N., Imms, F.J., Johnson, A.L., 1977. Falls in the elderly related to postural imbalance. BMJ 1, 261–264.

Pan, H.F., Hsu, H.C., Chang, W.N., Renn, J.H., Wu, H.W., 2016. Strategies for obstacle crossing in older adults with high and low risk of falling. J. Phys. Ther. Sci. 28, 1614–1620.

Pfister, A., West, A.M., Bronner, S., Noah, J.A., 2014. Comparative abilities of Microsoft Kinect and Vicon 3D motion capture for gait analysis. J. Med. Eng. Technol. 38, 274–280.

Rietdyk, S., Rhea, C.K., 2011. The effect of the visual characteristics of obstacles on risk of tripping and gait parameters during locomotion. Ophthalmic Physiol. Opt. 31, 302–310.

Sakurai, R., Fujiwara, Y., Ishihara, M., Higuchi, T., Uchida, H., Imanaka, K., 2013. Age-related self-overestimation of step-over ability in healthy older adults and its relationship to fall risk. BMC Geriatr. 13, 15–17.

Seo, N.J., Fathi, M.F., Hur, P., Crocher, V., 2016. Modifying Kinect placement to improve upper limb joint angle measurement accuracy. J. Hand Ther. 29, 465–473.

Tanaka, R., Takimoto, H., Yamasaki, T., Higashi, A., 2018. Validity of time series kinematical data as measured by a markerless motion capture system on a flatland for gait assessment. J. Biomech. 71, 281–285.

Tinetti, M.E., Speechley, M., Ginter, S.F., 1988. Risk Factors for Falls among Elderly Persons Living in the Community. N. Engl. J. Med. 319, 1701–1707.

Tomar, U.S., Gupta, N., 2012. An observational study of foot lifts asymmetry during obstacle avoidance. J. Neurosci. Rural Pract. 03, 324–327.

Wang, Q., Kurillo, G., Ofli, F., Bajcsy, R., 2015. Evaluation of Pose Tracking Accuracy in the First and Second Generations of Microsoft Kinect, in: 2015 International Conference on Healthcare Informatics. IEEE, pp. 380–389.

Xu, X., McGorry, R.W., Chou, L., Lin, J., Chang, C., 2015. Accuracy of the Microsoft Kinect™ for measuring gait parameters during treadmill walking. Gait Posture 42, 145–151.

Yen, H.C., Chen, H.L., Liu, M.W., Liu, H.C., Lu, T.W., 2009. Age effects on the inter-joint coordination during obstacle-crossing. J. Biomech. 42, 2501–2506.

Yeung, L.F., Yang, Z., Cheng, K.C.C., Du, D., Tong, R.K.Y., 2021. Effects of camera viewing angles on tracking kinematic gait patterns using Azure Kinect, Kinect v2 and Orbbec Astra Pro v2. Gait Posture 87, 19–26.

